# PMGen: From Peptide-MHC Structure Prediction to Peptide Generation

**DOI:** 10.1101/2025.11.14.688404

**Authors:** Amir H. Asgary, Amirreza Aleyasin, Jonas A. Mehl, Salman Fallah, Hasmig Aintablian, Burkhard Ludewig, Michele Mishto, Juliane Liepe, Johannes Soeding

## Abstract

Accurate structural modeling of peptide–MHC (pMHC) complexes is a prerequisite for understanding adaptive immunity and developing data-driven immunotherapies. However, current tools are often limited by narrow class coverage, restricted peptide lengths, or insufficient accuracy for downstream design tasks. Here, we introduce PMGen (Peptide MHC Generator), an integrated framework for structure prediction and structure-guided design of variable-length peptides across MHC class I and II. By introducing Initial Guess and Template Engineering as strategies to enforce anchor constraints in AlphaFold2, PMGen achieves state-of-the-art structural fidelity with median peptide core RMSDs of 0.54 Å for MHC-I and 0.33 Å for MHC-II, outperforming five state-of-the-art methods. We further demonstrate that PMGen captures the subtle structural impact of single-point neoantigen mutations and that model confidence (pLDDT) reliably correlates with structural accuracy. We investigated two potential applications of our framework: structure-aware peptide design and generating data for machine learning (ML) models. To this end, we introduced a framework to sample peptides with preserved structures and improved binding affinity. As an example for ML application, we fine-tuned ProteinMPNN on PMGen-modeled structures. This improved sequence recovery from 0.19 to 0.40 compared to the baseline. Ultimately, PMGen bridges the gap between high-fidelity structural prediction and downstream sequence design, offering a scalable solution to generate the large-scale, high-quality structural datasets required to train advanced predictive models in immunology. Available at https://github.com/soedinglab/PMGen.

## 1 Introduction

Major Histocompatibility Complexes (MHCs) are central to adaptive immunity. They present intracellular (MHC-I) or extracellular (MHC-II) peptides to T cells to trigger specific immune responses [1]. Antigenic peptides are vital for orchestrating immune activity, and their generation and presentation pathways are tightly regulated to ensure specificity and prevent autoimmunity. This regulation involves the generation of peptide antigens, transport and selective peptide binding to MHC molecules, activation of T cell receptors (TCRs), and modulation by regulatory T cells (Tregs). When this balance is disrupted, such as when self-peptides are misidentified as foreign, autoimmune diseases can result [2].

Designing peptides that bind effectively to MHC molecules has direct applications for immunotherapy and vaccine development. Identifying or engineering antigenic peptides with high binding affinity to specific MHC alleles enables the development of personalized treatments for cancer and infectious diseases. Immune checkpoint therapies, for example, have shown remarkable success in certain cancer patients by leveraging MHC-I-restricted neoantigen responses mediated by CD8^+^ T cells [3–9], and CD4^+^ T cell activation is at the cutting edge of targeted anti-cancer immunotherapies [10–12].

While tumors naturally present mutated neoepitopes, many are ineffective targets due to poor processing or weak MHC binding. In such cases, in silico redesign can improve immunogenicity while preserving structural similarity to the native antigen. A notable example is the engineered T210M variant of the melanoma-associated gp100 epitope, designed to enhance binding to HLA-A*02:01 [13]. To automate this discovery, reinforcement learning (RL)–based generators such as PepPPO [14] and RLpMIEC [15] have been proposed. These approaches, alongside established sequence-based predictors like NetMHCpan [16, 17] and others [18–21], learn from large-scale pMHC binding data. However, their utility in personalized medicine is often constrained by the inherent bias and class imbalance of training datasets; they perform well on common alleles but frequently fail to generalize to rare or novel MHC alleles where data are scarce.

Structural modeling offers a pathway to overcome these sequence-based limitations by leveraging the conserved architecture of the MHC binding groove. Both MHC classes share a similar fold: a *β* -sheet base flanked by two *α*-helices, with the peptide nestled between them [22]. Peptide binding is primarily driven by “anchor residues” that interact tightly with MHC pockets. Classical anchors include P2 and PΩ for MHC-I [23, 24], and P1, P4, P6, and P9 for MHC-II [25], with each allele having a preferred range of peptide lengths [26]. Accurate modeling of these interactions is essential to study the peptide mutation impact on T cell recognition [27]. Furthermore, restricting modeling to canonical anchor positions has been shown to significantly reduce peptide-core RMSD [28]. Recent work has demonstrated that such structural representations can be effective for sampling peptides under constraints beyond the MHC alone, including the TCR context [29]. Consequently, accurate pMHC structure prediction could provide the consistent training signals needed to build transferable models that generalize across MHC alleles and classes.

Despite this potential, accurate pMHC modeling remains a computational bottleneck. MHC-II modeling is particularly complex due to its dual-chain architecture and variable peptide length (11–25 residues), while MHC-I modeling struggles with predicting the flexible central core of the peptide [30]. Although AlphaFold [31] excels at protein folding, it often fails to correctly dock bound peptides [32]. Recent methods like Tfold [33], MHC-Fine [34], and PANDORA [28] attempt to address this by incorporating anchor information or fine-tuning. However, these tools face limitations such as restricted peptide length coverage, lack of MHC-II support, or high computational costs that hinder large-scale screening.

To address these limitations, we introduce Peptide MHC Generator (PMGen; https://github.com/soedinglab/PMGen), a unified pipeline for pMHC modeling and peptide design across both MHC classes. PMGen integrates anchor-guided AlphaFold modeling with structure-aware peptide generation. In benchmark evaluations, PMGen outperformed current state-of-the-art structure predictors. By bridging the gap between sequence-based prediction and structure-based optimization, PMGen provides a versatile platform for immunotherapy design, neoantigen modeling, and structural immunology.

## 2 Results

### PMGen Pipeline

PMGen consists of three core components: (1) the anchor feeding module, (2) structure prediction, and (3) peptide generation/binder selection (Figure 1A). The anchor feeding module takes peptide and MHC sequences as input, predicts anchor residues with NetMHCpan [16], and provides these as constraints to AlphaFold2 [35]. Alternatively, anchors can be specified by the user or selected based on the highest pLDDT score. In the structure prediction module We implemented two alternative approaches for incorporating anchor information into AlphaFold. First, Initial Guess (IG), which initializes AlphaFold2’s structure module with peptide anchor positions derived from templates with aligned anchors. Second, Template Engineering (TE), which performs anchor-constrained homology modeling using PANDORA, where the generated homology models are supplied to AlphaFold’s template module in place of real protein structures (see Methods). For peptide generation (Figure 1B), PMGen employs ProteinMPNN [36] to sample alternative peptides that align with the predicted backbone, either in a single step or iteratively. The sampled peptides can then be selected based on their predicted binding affinity to the MHC.

**Figure 1.**
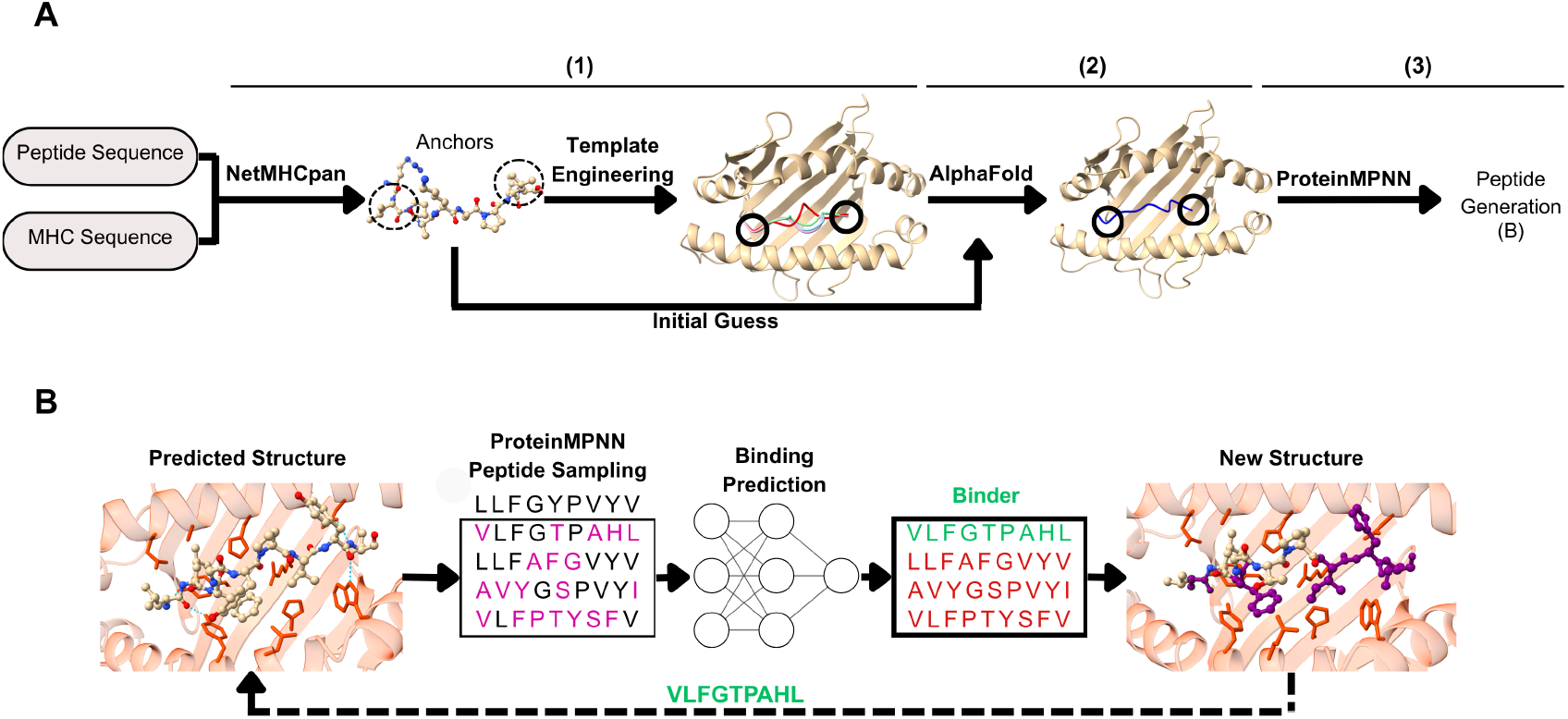
Anchor-guided modeling and structure-based peptide generation in PMGen. (A) For a given peptide–MHC pair, PMGen predicts anchor positions using NetMHCpan. Initial structural coordinates (Initial Guess) or Engineered Templates (with peptides from different templates shown in color) are used to input anchor position information to AlphaFold (1). AlphaFold performs anchor-aware structure prediction (2). The resulting backbone serves as input for ProteinMPNN to design alternative peptides (3). (B) Multiple peptide sequences are sampled using ProteinMPNN, and their binding affinities are predicted. The top-scoring peptide is then used for another round of structure prediction. This process can be repeated iteratively. Dashed lines represent the iterative loop, aiming to converge toward a high-affinity binder with an optimized sequence.

### Peptide-MHC Modeling Benchmark

To assess PMGen’s modeling performance, we compared it against five state-of-the-art tools: PANDORA [28], Tfold [33], AlphaFold Multimer [37], AFfine [38], and MHC-Fine [34]. We benchmarked PMGen in three modes: in its default Initial Guess mode (PMGen), in Template Engineering mode (PMGen+TE), and in an anchor-blind mode (PMGen+pLDDT). In the latter, the structure is selected based on the highest pLDDT score among

PMGen significantly outperformed all other methods across both MHC classes, achieving a median peptide-C*α*-RMSD (pRMSD) of 0.54 Å on MHC-I and 0.33 Å on MHC-II (Figure 2A). PMGen+pLDDT ranked as the second-best method, with median pRMSDs of 0.57 Å on MHC-I and 0.41 Å on MHC-II. Although PMGen+TE was less accurate than the IG mode, it still outperformed PANDORA, AFfine, and AlphaFold Multimer and achieved a performance comparable to Tfold, with a median pRMSDs of 1.04 Å on MHC-I and 0.49 Å on MHC-II.

**Figure 2.**
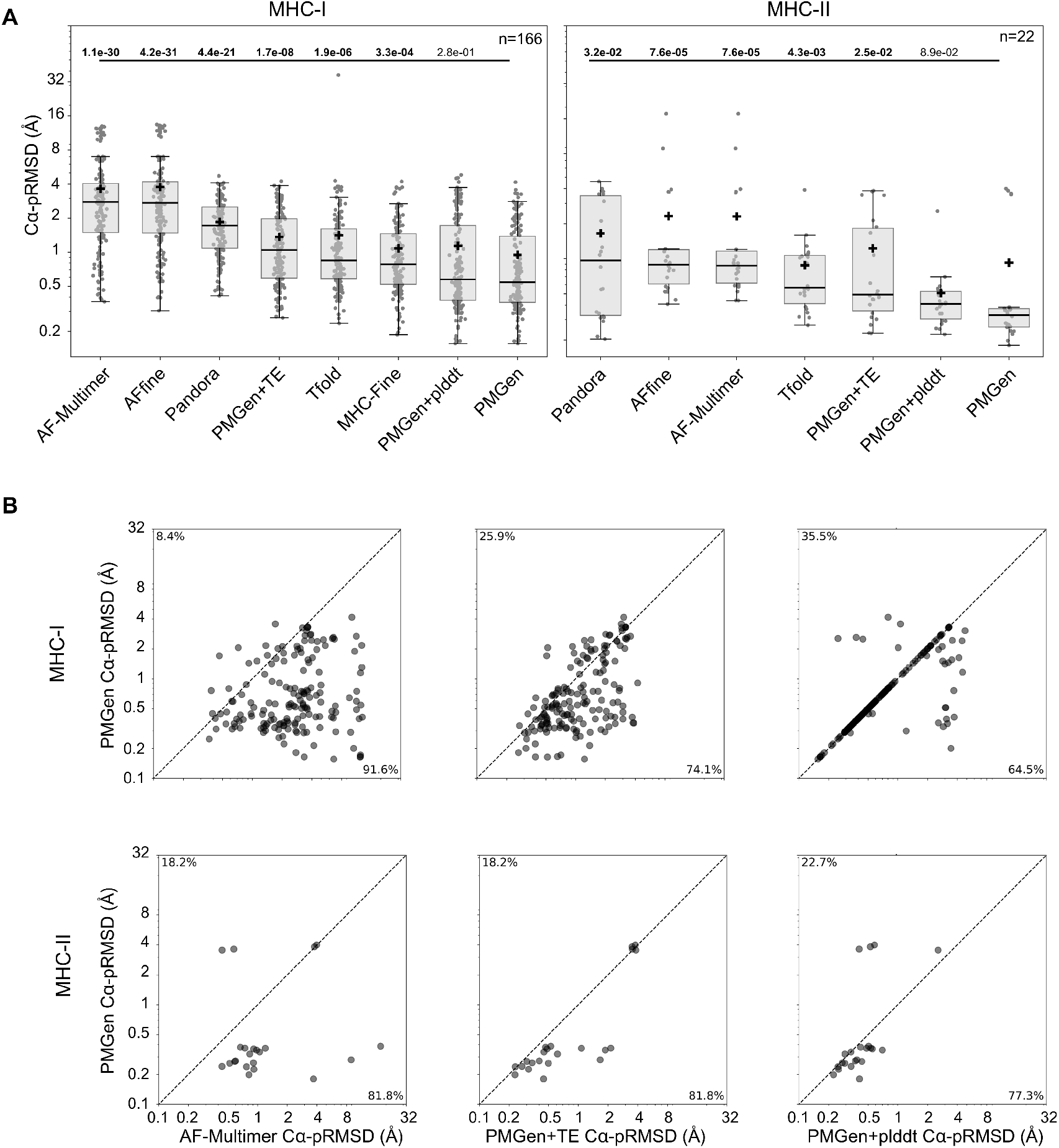
Comparative modeling assessment. (A) Modeling performance measured by RMSD over peptide C_*α*_ atoms. Left: MHC-I; right: MHC-II. Medians and means are indicated by **‘** −**‘** and **‘+’**, respectively. P-values from Wilcoxon signed-rank tests comparing each method with PMGen are shown above. Significant P-values (*P* < 0.05) are shown in bold. PMGen: using Initial Guess, PMGen+TE: using Template Enginneering, PMGen+pLDDT: anchor selection by pLDDT. (B) Pairwise comparisons of peptide C_*α*_ RMSD between PMGen and other methods. From left to right: AF-Multimer v2.2, PMGen+TE, and PMGen+pLDDT. Top row: MHC-I; bottom row: MHC-II. multiple initial guess runs over all possible anchor combinations (see Supplementary Methods). To exclude any structures used in AlphaFold2 training, we tested only on structures released after the cut-off date of 2018-04-30. Since AlphaFold3 [31] and the latest version of AF-Multimer have more recent training cut-offs, we utilized AF-Multimer v2.2 for benchmarking. We used AlphaFold model 2 ptm parameters for PMGen predictions due to its higher average pLDDT (Figure S2) on pMHC structures released before the benchmarking cut-off date (see Methods).

Pairwise pRMSD comparisons between PMGen and AlphaFold-Multimer v2.2 demonstrate that incorporating anchor information substantially improves structure prediction in nearly all cases (Figure 2B, left). This superior performance extends to comparisons against all other methods, with improvements observed in more than two-thirds of cases across both MHC classes (Figure S3). When comparing PMGen variants, the Initial Guess mode outperforms Template Engineering in the majority of cases (Figure 2B, middle). Interestingly, exhaustively testing all anchor combinations in IG mode and selecting the best structure based on pLDDT (PMGen+pLDDT) performs almost as well as direct anchor prediction, particularly for MHC-I (Figure 2B, right). This performance is supported by the high correlation of AlphaFold’s confidence metrics with pRMSD (Figure S4).

### Importance of Anchor Prediction in Structure Prediction

We evaluated the impact of anchor residue prediction on structural accuracy and assessed the concordance between the final structural anchor positions and those predicted by NetMHCpan. Previous work [27] characterized anchor positioning profiles, demonstrating that peptide residues with the lowest solvent-accessible surface area (SASA) strongly correlate with anchor function. Guided by this insight, we identified anchor positions for both ground-truth and predicted structures by selecting residues with minimal normalized SASA values and applying distance-based criteria (see Supplementary Methods). We then performed three comparisons:

1. NetMHCpan-predicted anchors *vs*. structurally defined anchors from predicted structures,
2. NetMHCpan-predicted anchors *vs*. structurally defined anchors from ground-truth structures, and
3. Structurally defined anchors from predicted structures *vs*. structurally defined anchors from ground-truth structures.

These comparisons were conducted using the benchmark PDB dataset. PMGen and PMGen+TE, which rely on NetMHCpan input, exhibited high agreement with NetMHCpan regarding the positioning of all four MHC-II anchors. As expected, PMGen+pLDDT, which operates without prior anchor constraints, positioned anchors differently. For MHC-II, agreement metrics were identical across all anchor positions (a1–a4) because the relative inter-anchor spacing is fixed; thus, a mismatch in the first anchor propagates to all subsequent positions (Figure 3A-right). In contrast, for MHC-I, all PMGen variants showed lower agreement with NetMHCpan at the first anchor position, while the second anchor exhibited near-complete concordance (Figure 3A-left).

**Figure 3.**
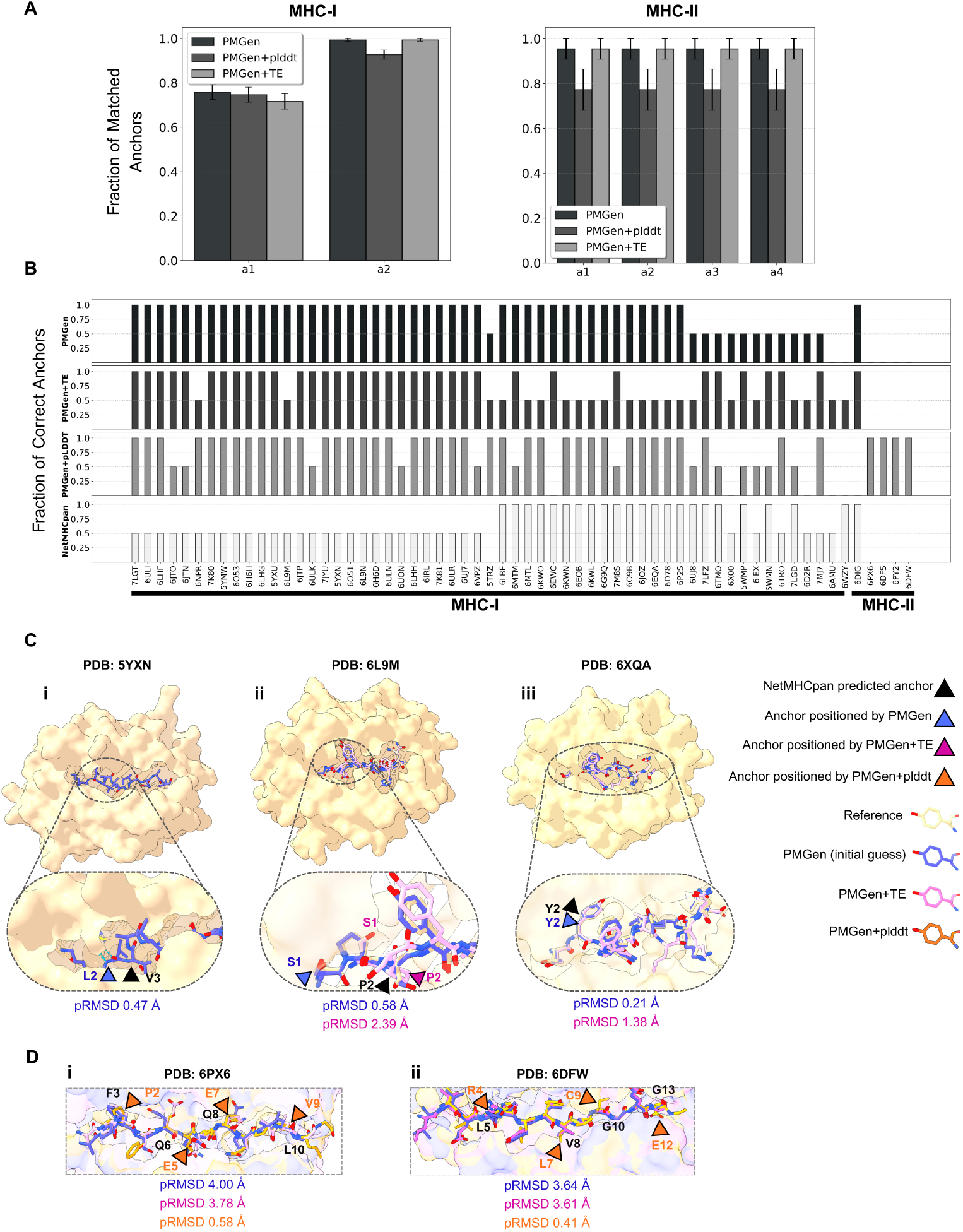
Anchor residue positioning analysis. (A) Fraction of anchors positioned that match the anchor positions predicted by NetMHCpan. (B) Structures in which at least one version of PMGen positioned an anchor differently from at least one anchor predicted by NetMHCpan. The Y-axis shows the fraction of correctly positioned/predicted anchors relative to ground-truth structures. (C) Examples of PMGen structure predictions where: **i**. NetMHCpan predicted the wrong anchor, but PMGen (IG) positioned it correctly; **ii**. PMGen+TE agreed with the incorrectly predicted anchors from NetMHCpan, while PMGen (IG) positioned the anchor correctly; **iii**. PMGen+TE, PMGen, and NetMHCpan all correctly defined the anchor, but the core region in PMGen+TE was wrongly predicted due to the influence of engineered templates. (D) PMGen+pLDDT correctly identified the anchors using only pLDDT scores, whereas PMGen (IG) and PMGen+TE misplaced the anchors due to incorrect predictions from NetMHCpan. pRMSD values and amino acid identities are shown for PMGen (blue), PMGen+TE (pink), PMGen+pLDDT (orange), the reference structure (tan), and NetMHCpan (black).

We identified 57 pMHC-I and 5 pMHC-II structures where at least one PMGen variant mismatched the ground-truth anchor positions. PMGen incorrectly positioned anchors that were correctly predicted by NetMHCpan in only five cases. Conversely, in the majority of instances, PMGen successfully corrected anchors mispredicted by NetMHCpan. Notably, for MHC-I, if NetMHCpan correctly predicted at least one anchor (typically anchor 2), PMGen generally retained this correct positioning. While PMGen+TE followed a similar trend, it occasionally misplaced anchors even when NetMHCpan predictions were correct—cases where PMGen’s IG mode maintained accuracy. Furthermore, PMGen+pLDDT correctly recovered anchor positions in 24 cases where NetMHCpan failed, though it introduced errors in 14 other cases (Figure 3B).

We further observed that the correction of mispredicted anchors by PMGen was associated with an improvement in pRMSD (Figure 3C-i). In contrast, the reduced performance of PMGen+TE appeared to arise not only from the mispositioning of anchors (Figure 3C-ii) but also from errors in the core conformation introduced by the engineered templates (Figure 3C-iii). For MHC-II, neither PMGen nor PMGen+TE could correct NetMHCpan mispredictions. However, PMGen+pLDDT successfully identified correct anchor positions in several instances where other variants failed (Figure 3B–D).

### Applications of PMGen in Structure-Aware Peptide Design and Model Fine-Tuning

As a secondary objective, we investigated PMGen’s applications in two fields: peptide design and generating accurate structures for training machine learning models. For the first purpose, we integrated ProteinMPNN [36] into the PMGen pipeline to enable structure-aware peptide generation for specific target MHC molecules (see Supplementary Methods). For the second purpose, we fine-tuned ProteinMPNN on predicted structures from an IEDB-derived pMHC dataset to improve ProteinMPNN’s main objective (sequence recovery) and to enhance higher-affinity peptide sampling.

For the first objective, we sampled peptides with protein MPNN conditioning on PMGen-predicted pMHC backbone, the corresponding MHC sequence, and any fixed (non-variable) peptide positions. We utilized the same PDB dataset from the structure prediction benchmark for evaluation. PMGen predictions were categorized into 51 high pRMSD (*>* 1.0 Å) and 55 low pRMSD (*<* 0.6 Å) structures. We performed three in silico screens with sliding windows of single, double, and triple consecutive amino acid substitutions. For each window, we generated peptide variants using ProteinMPNN and selected candidates based on their NetMHCpan-predicted %EL ranks. Randomly mutated peptides served as controls (see Supplementary Methods for sampling details). After predicting the structures of sampled and randomly mutated peptides, we calculated the pRMSDs vs the original peptide’s predicted structures. Cumulative enrichment analyses were conducted for each screen and structure separately (Figure S5). We measured the enrichment of ProteinMPNN-sampled peptides relative to the random mutation baseline in terms of pRMSD (Figure 4A). The results demonstrated higher enrichment (AUC) for PMGen predictions with low pRMSD (Figure 4B). The absolute pRMSD comparison between ProteinMPNN-sampled peptides across good and bad predictions indicates lower deviation from the original structure in the good prediction group (Figure 4C). These results indicate that PMGen high-quality structures can be used to sample peptides with similar structural properties.

**Figure 4.**
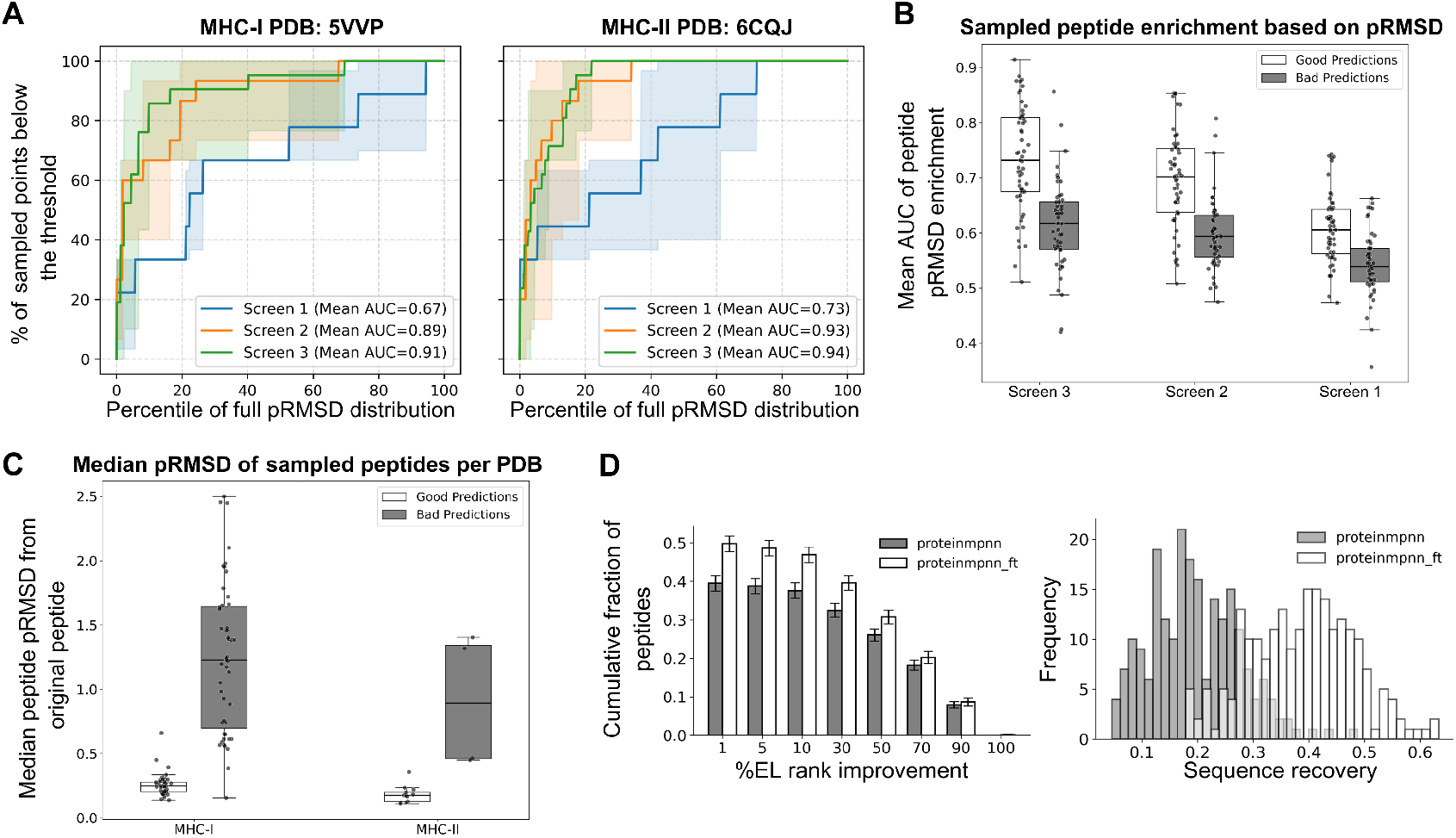
Peptide generation analysis conditioned on backbone and MHC sequence. (A) Cumulative enrichment analysis of peptide RMSD (pRMSD) for PMGen-generated peptides compared to randomly mutated peptides in two sample PDBs. Screens 1, 2, and 3 correspond to sliding windows with one, two, and three point mutations, respectively. A higher AUC indicates greater enrichment among peptides with lower pRMSD. (B) Comparison of pRMSD enrichment AUCs between good (low original pRMSD, *n* = 55) and bad (high original pRMSD, *n* = 51) predicted structures. (C) Median pRMSD values of generated peptides across all screens; each point represents the median deviation of generated peptides from the original predicted structure for a single PDB. (D) Cumulative fraction of sampled peptides with improved (lower) EL percentile rank compared to the original peptide (left) and sequence similarity distribution of sampled peptides to the original peptide (right). Peptides were generated from the same predicted structure using the original ProteinMPNN and the version fine-tuned on PMGen structures.

For the fine-tuning application, we predicted 10,216 high-confidence pMHC structures using PMGen, filtered by pLDDT (≥ 80) and peptide PAE (≤ 6.0), spanning 297 MHC-I alleles (see Supplementary Methods and Figure S6A-B). We fine-tuned ProteinMPNN using sequence recovery, consistent with its original training objective (Figure S6C). Fine-tuning increased peptide sequence recovery on the test set from a mean of 0.19 to 0.40 (Figure 4D, right). We also observed that at least 50% of sampled peptides from the fine-tuned model had higher predicted affinity than the original peptide, compared to 40% for the baseline ProteinMPNN (Figure 4D, left).

### PMGen Evaluation on a Neoantigen/Wild-Type Pair Test Case

To demonstrate PMGen’s capability in distinguishing subtle structural differences between antigens, we evaluated its performance on a wild-type/neoantigen pair previously used to benchmark Tfold [33]. Specifically, we modeled the wild-type antigen KLSHQ**P**VLL bound to HLA-A*02:01 (PDB: 8TBW) and its single-point mutation neoantigen variant KLSHQ**L**VLL (PDB: 8U9G). PMGen was executed in initial guess mode for both cases, and the predicted models were compared to the experimental X-ray structures.

For the wild-type complex, PMGen achieved a pRMSD of 0.80 Å (Figure 5A), a notable improvement over the previously reported Tfold pRMSD of 1.2 Å. The predicted peptide orientation and anchor positioning closely resembled the crystal structure (Figure 5A-i). Residues K1, L2, S3, L8, and L9 were closely aligned with the experimental coordinates, while P6 and V7 showed a minor shift in their C*α* positions (Figure 5A-ii). Side-chain rotamers for H4 and Q5 showed minor deviations, though the backbone alignment was largely preserved.

**Figure 5.**
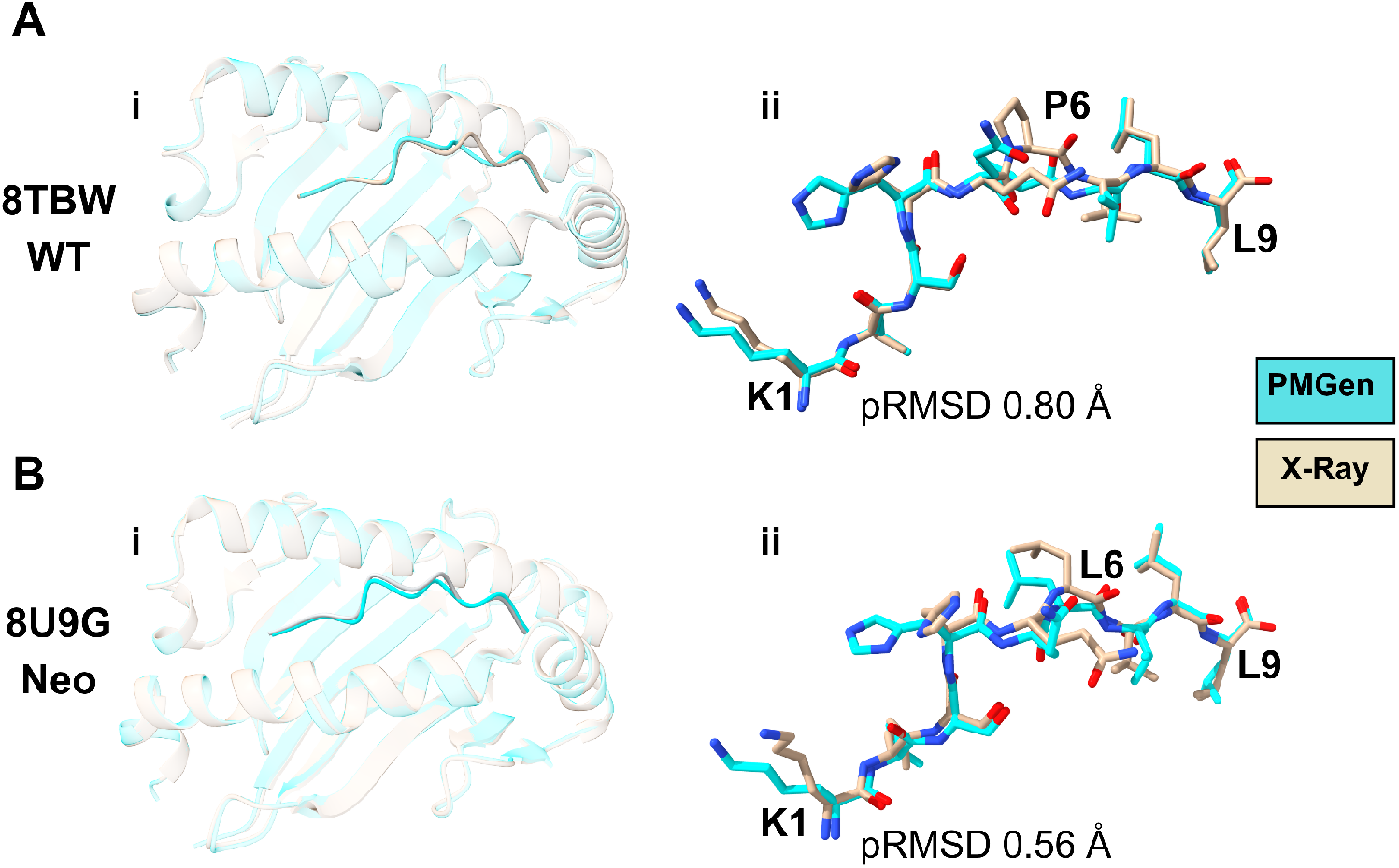
Wild-type antigen vs. neoantigen modeling with PMGen. (A) Wild-type antigen KLSHQPVLL: (i) Predicted structure (cyan) aligned with the X-ray structure (amber). (ii) Peptide alignment showing that C*α* atoms of almost all residues match, with the exception of P6. (B) Neoantigen KLSHQLVLL: (i) Predicted structure (cyan) aligned with the X-ray structure (amber). (ii) PMGen correctly predicts the side chain of L6 pointing outwards.

For the neoantigen (P6→L6), PMGen achieved a pRMSD of 0.56 Å (Figure 5B-i), performing comparably to Tfold (0.6 Å), with C*α* positions closely aligned to the reference structure. In the experimental structure, the side chain of the mutated L6 is oriented outward and slightly toward an MHC *α*-helix. PMGen correctly predicted this orientation. Similar to the wild-type case, the predicted rotamers for H4 and Q5 differed slightly from the experimental structure (Figure 5B-ii).

## 3 Methods

### pMHC PDB Data Processing

We collected PDB files containing bound-state peptide–MHC structures from the IMGT [39] and RCSB Protein Data Bank [40] databases. Structures with a reported resolution worse than 3.5Å were excluded. All PDB files were cleaned and renumbered, and non-amino acid molecules were removed using the Biopython PDB module [41].

For structural alignment, each chain of the collected PDB structures was aligned against the corresponding chain of selected reference structures: 4H25 for MHC-II and 4U6Y for MHC-I. Immunoglobulin-like domains (*β* 2 and *α*2 for MHC-II; *α*3 for MHC-I) were manually removed from the references, retaining only the peptide-binding domains. Structural alignment was performed exclusively on these binding domains using TMalign [42], after which all non-aligned MHC regions were discarded.

In cases where PDB files contained additional chains (e.g., TCRs), only the peptide and MHC chains were retained. The peptide chain was identified as the chain with length ≤ 25 amino acids having the smallest Euclidean distance to the center of mass (COM) of the MHC binding domain.

This preprocessing yielded 964 structures (116 MHC-II and 848 MHC-I). We subsequently excluded 40 structures due to peptide lengths *<* 8 amino acids or processing failures in at least one benchmarking tool (Table S1). For benchmarking, we selected only structures deposited after the AlphaFold 2.2 training cut-off date of 30 April 2018, resulting in a final test set of 22 pMHC-II and 166 pMHC-I structures. The remaining pre-cut-off data served as a discovery set for optimizing AlphaFold model parameters (based on overall pLDDT) for use in PMGen (Figure S2).

### Template Engineering

PMGen takes peptide and MHC sequences as input to predict the pMHC structure and binding affinity, with an option for peptide generation. The workflow begins by identifying potential peptide anchor positions, which are either predicted by NetMHCpan or specified by the user. For long peptides with multiple predicted anchors, users can retain the top *k* anchors ranked by NetMHCpan’s %EL and binding affinity (see Supplementary Methods). In this study, we used only the top-ranked anchor for benchmarking.

To identify the MHC allele, PMGen aligns the query MHC sequence against a local repository of MHC sequences with known structures, selecting the closest match by sequence similarity. Alternatively, the allele can be user-specified. If the identified allele is present in NetMHCpan’s supported list, it is used directly for anchor prediction. Otherwise, PMGen selects the most similar supported allele based on alignment score solely for anchor prediction, preserving the original query sequence for modeling.

The query sequences and defined anchor positions are then processed by a modified PANDORA pipeline [28] (see Supplementary Methods). PANDORA performs a BLAST search against the local pMHC database to select the top homologous structure, identifies the MHC anchor-binding pockets, and uses the coordinates of the peptide’s anchor residues as spatial constraints. Anchor-constrained homology modeling is subsequently performed using MODELLER [43].

We term these constrained structures *engineered templates*. While the peptide geometry within the MHC binding groove (anchors) remains consistent across templates, the core and flanking regions vary. PMGen generates multiple engineered templates per query (default: four), ranking them by MODELLER’s molpdf score, and supplies them as input templates to AlphaFold.

### AlphaFold Initial Guess Implementation

To reduce computational overhead, we implemented an “Initial Guess” (IG) mode inspired by Bennett et al. [44], which bypasses the template engineering step. We designated this as the default mode for PMGen.

In IG mode, peptide anchor positions are predicted (see Supplementary Methods) or user-specified. A BLAST search is performed against the local pMHC PDB database (see Template Engineering in Methods), and the *k* structures with the highest alignment scores are selected as templates.

A custom alignment file is generated containing paired MHC–peptide alignments for each selected template. Crucially, peptide anchor positions are aligned first to ensure accurate transfer, followed by the remaining residues. The aligned templates are then provided to AlphaFold. The 3D coordinates of the peptide anchors and all MHC residues are supplied directly to AlphaFold’s structural module. In contrast, the coordinates of the remaining (core) peptide residues are masked. This effectively compels AlphaFold to model these positions flexibly during its first structure-prediction iteration, prior to standard recycling refinement.

## 4 Discussion

PMGen is a versatile framework for pMHC structure prediction and structure-guided peptide design, integrating anchor residue information into AlphaFold2 [35] via two modes: Initial Guess and Template Engineering. Unlike existing tools such as MHC-Fine [34], which are limited to MHC-I and short peptides, PMGen supports both MHC classes and a broad range of peptide lengths. Benchmarking against state-of-the-art methods [28, 33, 34, 37, 45] demonstrated PMGen’s superior structure prediction accuracy, with a pRMSD median of 0.54 Å for MHC-I and 0.33 Å for MHC-II (Figure 2A). The exclusion of overlapping templates and AlphaFold training structures during evaluation ensured unbiased performance comparisons, establishing PMGen as the top-performing pMHC structure prediction algorithm despite potential validation set overlap in competing tools.

Our results demonstrate that AlphaFold effectively assimilates anchor information from both IG and TE modes without requiring parameter fine-tuning (Figure 2B). This inherent flexibility aligns with recent findings that AlphaFold can leverage contact maps and distance constraints to resolve structural ambiguities [46, 47]. Furthermore, the utility of providing multiple structural priors to enhance prediction and design is well-documented [44, 48]. Remarkably, we observed that the Initial Guess strategy outperformed Template Engineering (Figure 2B, middle). We hypothesize that this improvement stems from how AlphaFold processes conflicting template data. When presented with multiple similar templates, the model tends to converge on an average consensus conformation with artificially high confidence. This behavior restricts the search space to the close neighborhood of the engineered templates. In contrast, the IG mode provides only coarse spatial constraints relative to the MHC. This looser guidance enables AlphaFold to explore a broader conformational landscape. Consequently, IG correctly resolved structures in challenging cases where the more restrictive TE mode misplaced the core region (Figure 3C).

PMGen profits from being provided with correct anchor positions (Figure 3A), yet it demonstrates a robust capacity to recover correct positions even when anchor inputs are wrong (Figure 3B). This highlights AlphaFold’s inherent denoising capability regarding partial conformations [49]. Since different PMGen modes yield distinct structural ensembles, their combination increases the probability of sampling the native conformation. Crucially, we observed that correct anchor positioning correlates strongly with higher pLDDT scores (Figure 2B, right). This relationship allows for effective model ranking (Figure S4), consistent with established correlations between pLDDT and pRMSD in pMHC complexes [33]. The practical value of this metric was evident in four challenging MHC-II cases (6PX6, 6DFS, 6PY2, 6DFW) where sequence-based methods failed. In these instances, prioritizing models based on pLDDT (PMGen+pLDDT) rather than predicted anchors successfully resolved the correct core (Figure 3B–D). These findings indicate that leveraging structural confidence scores can effectively mitigate errors in purely sequence-based anchor prediction.

PMGen’s accurate structures enable translational applications, such as structure-aware peptide generation to profile MHC binders and providing training data for structure-based machine learning models. To this end, we integrated ProteinMPNN [36] into our workflow to leverage its autoregressive sampling. We used NetMHCpan [16] to rank sampled candidates in the current paper. This choice was motivated by its broad allele coverage [50]. Our enrichment analysis revealed that generated peptides preserve the structure of the original peptide. This effect was especially pronounced in screens with a higher number of mutations (Figure 4A-B). We showed that high structure preservation is achieved with high-quality predicted structures (Figure 4C). As a second application, we showed that PMGen-predicted structures can be used to train machine learning models. In the current study, we fine-tuned ProteinMPNN as an example. This successfully improved peptide sequence recovery from 0.19 to 0.40 without compromising the original model weights for MHC sequence recovery (Figure S6, Figure 4D). As an indirect metric, we observed that upon fine-tuning the fraction of peptides with higher predicted affinity also increased (Figure 4D). While in the current study we focused on binding affinity, in silico metrics can be optimized for specific goals, such as TCR-specific rankings [51]. These findings emphasize the possible applications of PMGen high-quality predicted structures in vaccine design and machine learning fields.

Beyond these computational applications, PMGen’s structural accuracy also has direct implications for neoantigen discovery and optimization. Our method addresses critical limitations in current neoantigen prediction tools, which often rely on sequence-based features or low-resolution structural models [52, 53]. By providing high-quality 3D structures and enabling structure-aware generation, PMGen facilitates the rational design of immunogenic neoantigens [29] and enhanced mimotopes. In a benchmark case, PMGen accurately captured mutation-induced conformational changes that are critical for altered T-cell recognition (Figure 5) [54, 55]. Such mimotope-based strategies have shown promise in preclinical models and early-phase clinical trials [56–60]. They also offer similar therapeutic potential in autoimmune diseases [61]. PMGen can offer faster alternatives to MD-based frameworks by screening over the structure of thousands of candidate peptides [62]. Recent work demonstrates that incorporating 3D MHC-II coordinates yields experimentally validated neoantigens [55]. PMGen’s dual-class support extends this paradigm. This capability is particularly important, as combining MHC-I and MHC-II epitopes enhances therapeutic efficacy [63]. While the current study primarily establishes the PMGen framework, its direct application to clinical vaccine design remains a promising avenue for future investigation.

Several limitations of the current study should be acknowledged. First, PMGen relies on AlphaFold2. While our anchor-constraint strategy substantially improves peptide docking performance, newer architectures like AlphaFold3 [31] may offer further enhancements. However, AlphaFold3’s recent training cut-off complicates unbiased benchmarking against existing methods. Second, the default anchor prediction depends on NetMHCpan. This tool may underperform for rare or poorly characterized alleles, though the pLDDT-based selection mode provides a sequence-independent fallback. Third, our peptide generation module shows encouraging in silico enrichment for predicted binding affinity. However, these candidates have not yet been experimentally validated. Predicted binding affinity is a necessary but insufficient proxy for immunogenicity. This property additionally depends on antigen processing, peptide stability, and the TCR repertoire. Finally, the current design module optimizes for MHC binding without explicitly modeling TCR interactions. This constraint could be addressed through integration with emerging TCR-pMHC modeling approaches [48].

PMGen offers a robust and versatile framework for pMHC modeling and neoantigen design. The pipeline is well-suited for integration into both experimental workflows for cancer immunotherapy and autoimmune research, as well as structure-based in silico pipelines. By enabling the high-throughput generation of accurate structural models, PMGen serves as a critical first step in producing high-quality synthetic training data for machine learning frameworks, directly addressing the limitations and biases inherent in current experimental and computational datasets. Our future work will focus on the *in vitro* and *in vivo* validation of PMGen-derived neoantigens, improved model interpretability, and integration with advanced generative architectures [15, 64] to accelerate the development of next-generation immunotherapies.

## Key Points

- PMGen is a unified framework for peptide–MHC structure prediction across both MHC class I and II, outperforming five state-of-the-art methods with median peptide core RMSDs of 0.54 Å and 0.33 Å, respectively.
- Incorporating anchor residue constraints into AlphaFold2 via Initial Guess or Template Engineering substantially improves peptide docking accuracy without requiring model fine-tuning.
- AlphaFold2 confidence scores (pLDDT) reliably predict structural accuracy, enabling sequence-independent anchor selection that rescues cases where NetMHCpan anchor prediction fails.
- PMGen-guided ProteinMPNN sampling generates peptide variants with enriched MHC binding affinity while preserving structural similarity to the original peptide.
- High-quality pMHC structures predicted by PMGen provide a scalable source of synthetic structural data, with demonstrated utility for both training machine learning models and structure-aware peptide design.

## Abbreviations

(AF): AlphaFold
(AFfine): AlphaFold fine-tuned
(AUC): Area Under the Curve
(BLAST): Basic Local Alignment Search Tool
(COM): Center of Mass
(CD4+ T-cells): Cluster of Differentiation 4 Positive T-cells
(CD8+ T-cells): Cluster of Differentiation 8 Positive T-cells
(EL): Eluted Ligand
(HLA): Human Leukocyte Antigen
(IEDB): Immune Epitope Database
(IG): Initial Guess
(MHC): Major Histocompatibility Complex
(MHC-I): MHC Class I
(MHC-II): MHC Class II
(MSA): Multiple Sequence Alignment
(pMHC): peptide-MHC
(pRMSD): peptide C*α* Root Mean Square Deviation
(pLDDT): predicted Local Distance Difference Test
(PAE): Predicted Aligned Error
(PDB): Protein Data Bank
(Tregs): Regulatory T-cells
(RL): Reinforcement Learning
(RMSD): Root Mean Square Deviation
(SASA): Solvent Accessible Surface Area
(TCR): T-Cell Receptor
(TE): Template Engineering

## Ethics Statement

This study is based entirely on computational analysis of publicly available data from the Protein Data Bank and IEDB database. No ethical approval was required as no human subjects, human tissue, or animals were involved. The authors declare no competing interests.

## Data availability

The datasets used for benchmarking and fine-tuning are available from the IEDB database and PDB as described in the Methods. All the generated structures are available at https://wwwuser.gwdguser.de/~compbiol/pmgen/pdbs.

## Code availability

The source code for PMGen is available on GitHub https://github.com/soedinglab/PMGenrepository. The code and data to reproduce paper results is available at https://wwwuser.gwdguser.de/~compbiol/pmgen/. A user-friendly notebook to run PMGen is available at https://colab.research.google.com/github/soedinglab/PMGen/blob/master/colab.ipynb. The code developed to fine-tune ProteinMPNN, including model weights is available at https://github.com/AmirAsgary/ProteinMPNNFineTune.

