## Supplementary information for "PMGen: From Peptide-MHC Structure Prediction to Peptide Generation"

### Binding Groove Distance Distribution

We selected the longest sequences among the 964 processed structures (PDB ID: 4U6Y for MHC-I, 4H25 for MHC-II). We then manually defined the right, core, and left regions on the $\alpha$ structures. The right region was identified as the set of amino acids $\{4,5,6\}$ following the nearest MHC$_\alpha$ residue to the first tightly bound peptide amino acid (with a $C_\alpha$ distance $< 4.5$ Å). The left region was defined using the same rule but on the opposite side, primarily where long peptides protrude from MHC-II. The core region was considered a broader segment between the right and left regions, consisting of 10 amino acids. For each region, the nearest partner amino acids on the opposing MHC$_\alpha$ secondary structure were identified based on their sequence position and the middle amino acid's Euclidean distance. These distances were averaged for each side. For the remaining structures, we determined the right, core, and left regions by aligning their amino acids to those of the selected structures (Figure S1). Alignments were performed using Clustal Omega multiple sequence alignments (MSAs) [1].

Table S1: Excluded PDBs for benchmarking

| | | | | | | | | | |
|---|---|---|---|---|---|---|---|---|---|
| 3VFU | 1ED3 | 4NO0 | 3VFT | 3VFO | 3VXU | 2ICW | 7MJ8 | 5DDH | 4QRP |
| 4JRY | 6RP9 | 4JQX | 1ZHL | 6AT5 | 6LF9 | 3VFR | 4JRX | 4U6Y | 4ZUT |
| 4ZUV | 1ZHK | 3W0W | 4NO2 | 3VFP | 4ZUW | 3VFW | 3VFM | 3KXF | 3VFV |
| 1XH3 | 6LF8 | 3BW9 | 6NF7 | 2AK4 | 1R5I | 1ZT1 | 3VFN | 3VFS | 2OJE |

### AlphaFold Prediction

To predict final structures, we used the Python implementation of AlphaFold [2]. This implementation also provides a fine-tuned variant, AFfine, trained specifically for the pMHC classification task. Both implementations are based on the AlphaFold2 architecture and adapted for multi-chain prediction.

Multi-chain functionality is enabled by shifting the residue indices of each chain by 200. This prevents AlphaFold from interpreting separate chains as a continuous polypeptide. For each prediction, the query sequence was aligned to the engineered templates and supplied to AlphaFold.

The number of recycling iterations in AlphaFold was kept at the default value of three. For benchmarking, we used AlphaFold's `model_2_ptm` as well as the AFfine parameter set.

### Structure Prediction Benchmarking Setting

For benchmarking with AlphaFold-Multimer 2.2, we used a local installation of ColabFold v1.5.5 [3]. Multiple sequence alignments and template searches were performed against the UniRef30 and PDB70 databases, respectively. Predictions were generated using the five default AlphaFold models, and only the top-ranked structure reported by AlphaFold was used in the benchmarking analysis.

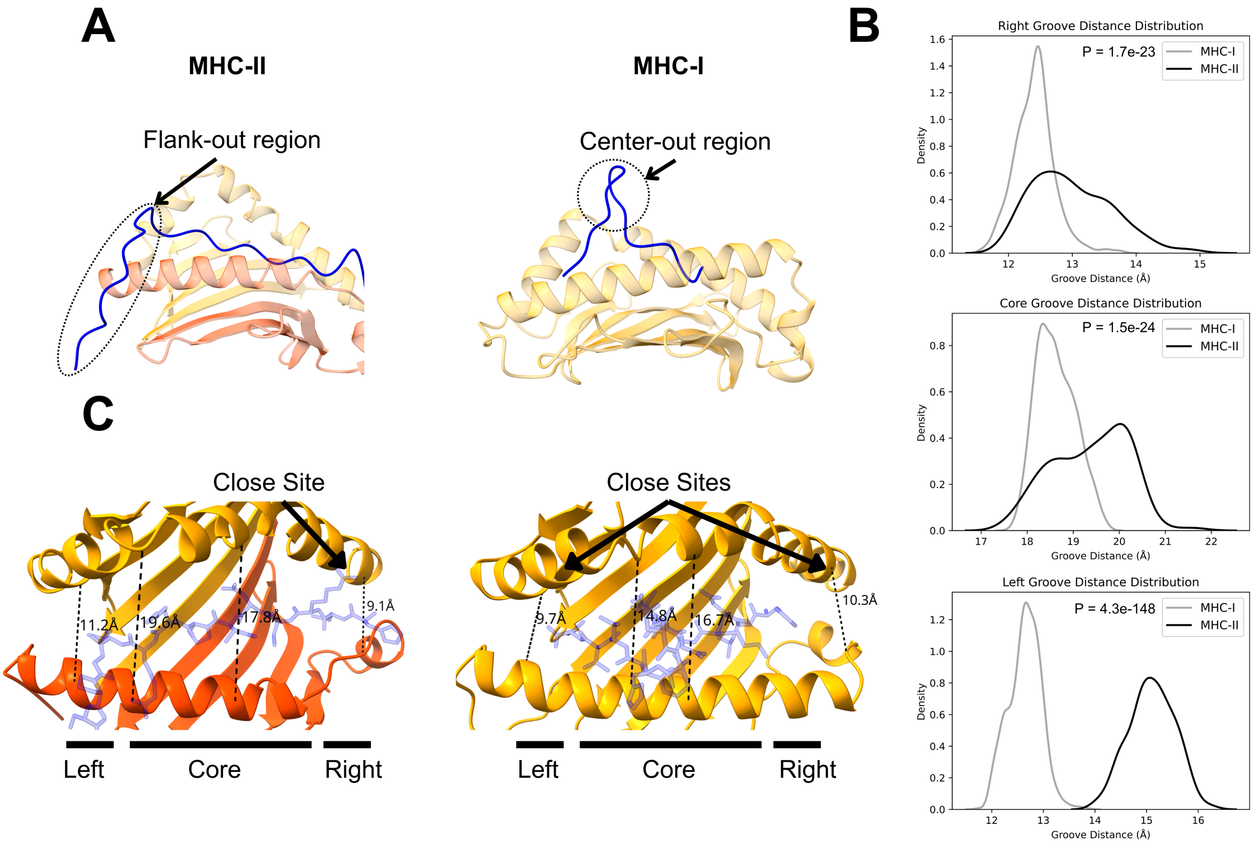


Figure S1: **Distinct binding properties of MHC-I and MHC-II.** (A) MHC-I tends to form a core-out conformation, whereas MHC-II adopts a flank-out conformation. (B) The binding groove of MHC-I has a smaller distance at both the right and left sites, while a larger central groove distance indicates the peptide's tendency to bulge outward. In contrast, MHC-II shows partial overlap with MHC-I in the right and core groove regions but exhibits complete separation in the left region due to its flank-out property. (C) Illustration of the binding groove regions in MHC structures: right (peptide entry site), core, and left (peptide exit site).

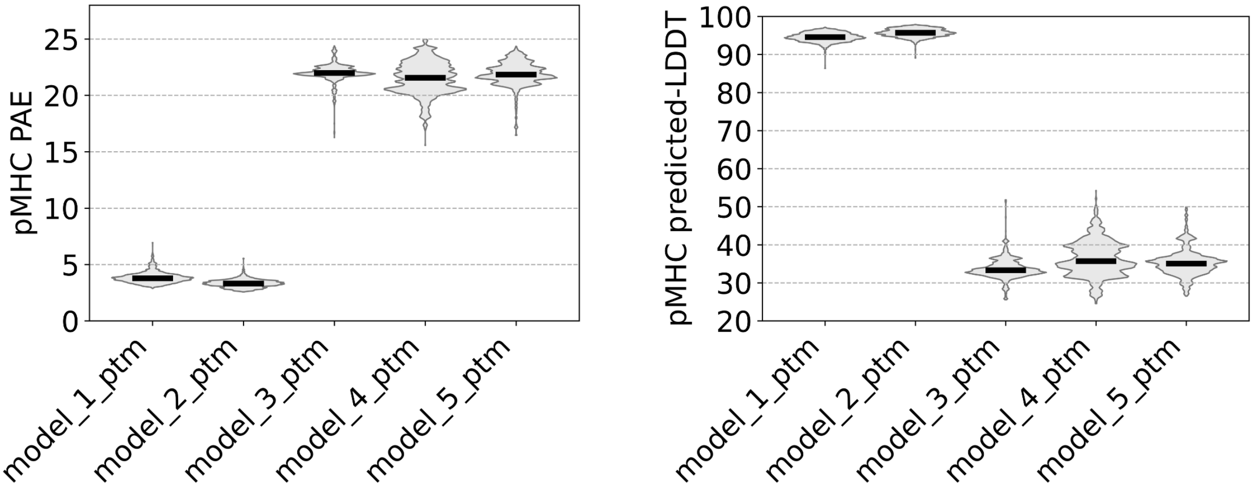


Figure S2: **AlphaFold2 performance.** The performance of different models in peptide-MHC structure prediction is visualized on the discovery set averaged over predictions on different numbers of templates (1 to 6).

For AFfine predictions, template PDB files were obtained from ColabFold search results. MSAs were omitted to maintain consistency with the original AFfine pipeline [2]. We employed AFfine's fine-tuned version of `model_2_ptm_ft`, which was trained on the classification task of peptide–MHC binding. We did not do the full alignment implementation explained in the paper, as our aim was to only test if fine-tuned weights can show superior performance than original weights.

Tfold [4] was executed with default settings, using its integrated SeqNN model for anchor prediction. As Tfold outputs multiple ranked structures, only the top-ranked structure generated by the tool was considered for benchmarking.

PANDORA [5] predictions were performed with default settings, using the built-in PANDORA BLAST and PDB databases for homology search and template selection. To automate anchor position assignment, we enabled the NetMHCpan option [6].

MHC-Fine [7] was executed locally with default settings. MSA generation involved jackhmmer queries to its dedicated database for MHC complexes.

For PMGen and PANDORA benchmarking, we applied a leave-one-out strategy during template search and engineering. Specifically, templates with the same PDB ID as the target structure were excluded to avoid potential bias.

For Multimer 2.2, Tfold, and MHC-Fine, the template search was performed internally by the respective tools, which do not implement a leave-one-out benchmarking protocol. Therefore, for these methods we relied solely on the training structure cut-off date to prevent information leakage, and identical template sequences were not explicitly excluded.

All tools were benchmarked on both MHC-I and MHC-II, except for MHC-Fine, which only supports MHC-I. To ensure fairness, deviations from the default tool settings were minimized. For AlphaFold-based methods, parameters with a training cut-off of 2018 were used by default. All tools were benchmarked on PDB structures released after this training cut-off.

### Structure Prediction Evaluation Metrics

To compare structure prediction models, the primary evaluation metric was the root-mean-square deviation (RMSD) of the C$\alpha$ atoms between the predicted and true structures of a peptide $i$ with length $L$, after superposition based on the MHC structures:

$$\mathrm{pRMSD}_i = \sqrt{\frac{1}{L}\sum_{j=1}^{L}\left\|\mathbf{r}_{i,j}^{\mathrm{CA,pred}} - \mathbf{r}_{i,j}^{\mathrm{CA,true}}\right\|^2}.$$

The predicted aligned error (PAE) was symmetrized for each residue pair using

$$\mathrm{PAE}_{ij}^{\mathrm{sym}} = \frac{\mathrm{PAE}_{ij} + \mathrm{PAE}_{ji}}{2}.$$

Only amino acid pairs within a 10 Å radius were considered when computing the final average per-residue PAE. This value is reported and used throughout the current study.

The predicted lDDT scores provided directly by AlphaFold 2 were also used as an evaluation metric.

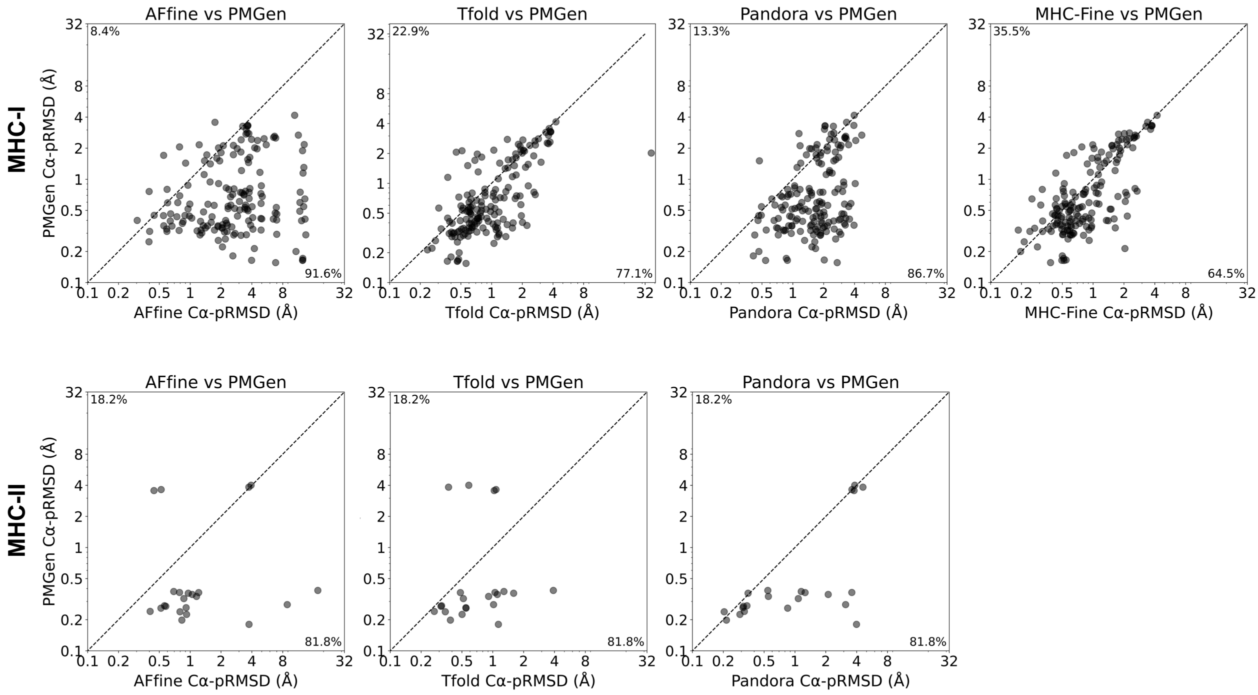


Figure S3: **Pairwise comparison between PMGen and other methods.**

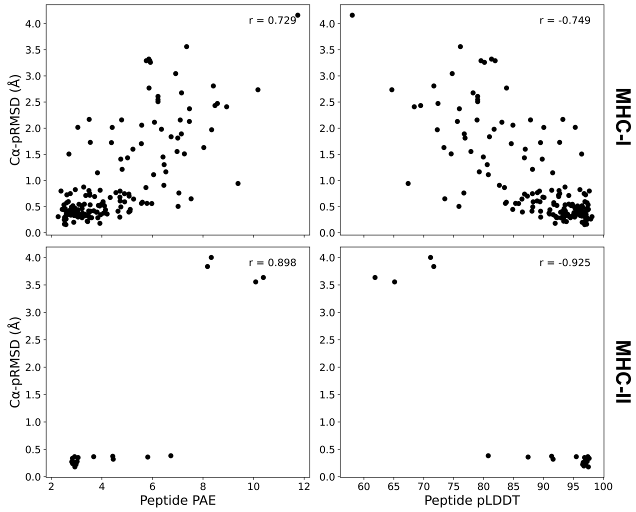


Figure S4: **PMGen C$\alpha$−pRMSD correlation with PAE and pLDDT.**

### Defining Anchor Residues from Structure

To identify anchor residues in each predicted pMHC structure, we used the solvent accessible surface area (SASA) as a selection criterion. SASA has been shown to be a reliable structural indicator for identifying peptide anchor positions [8]. We calculated the SASA for each residue as the total solvent-accessible surface area of its atoms, using the FreeSASA package [9]. Since amino acids inherently have different range of SASA values, we normalized the total surface area by maximum allowed solvent accessibilities for each amino acid [10].

Anchor positions were determined directly from the pdb structures rather than from sequence conservation, ensuring that the definition was minimally affected by sequence identity. Our method selects residues that (i) have low SASA values and (ii) satisfy a minimum separation in sequence index (difference between their positions in the peptide sequence).

Let $P$ be the set of all peptide residue positions in a given peptide–MHC complex. We define $F$ as the set of sets $S$ of all feasible anchor position combinations:

$$F = \left\{ S \subseteq P \,\middle|\, |S| = m, \forall p \in S : \quad SASA_p < 50, \quad \text{and} \quad \max(S) - \min(S) > n \right\}$$

The selected anchor residues $A$ are then given by:

$$A = \arg\min_{S \in F} \sum_{p \in S} SASA_p \tag{1}$$

$$\text{with} \quad \begin{cases} m = 2, & n = 6, \quad \text{if MHC-I} \\ m = 4, & n = 8, \quad \text{if MHC-II} \end{cases}$$

Here, $m$ denotes the total number of anchor residues to be selected, and $n$ is the minimum sequence index distance allowed between two consecutive anchors. For MHC-II, we also check if the space between each consecutive anchor is at least 2. The residues in $A$ are therefore those that satisfy the SASA and spacing constraints while minimizing the total SASA sum.

### Evaluation of Modeling Performance

To evaluate the accuracy of the predicted pMHC structures, we first aligned the MHC chain(s) of each predicted structure to the corresponding chain(s) in the ground-truth structure. The alignment was performed using only MHC chains so that the resulting superposition was dominated by MHC structure. For MHC-II structures, both chains were treated as a single concatenated chain during alignment to preserve their relative conformation.

The rotation matrix obtained from the MHC alignment was then applied to the peptide coordinates in the predicted structure, thereby placing the peptide in the same reference frame as the ground-truth complex.

The peptide core region was defined as the segment between the first and last anchor residues, where anchors were identified separately for both the ground-truth and predicted structures (see Supplementary Methods). As our primary structural accuracy metric, we computed the mean RMSD of the C$\alpha$ atoms within the peptide core region.

To assess the accuracy of anchor positioning, we compared the index positions of each corresponding anchor in the prediction and reference structures (e.g., anchor 1 index in ground-truth versus anchor 1 index in prediction). The fraction of matched anchors between NetMHCpan predicted anchors and predicted or ground-truth structures was calculated with the same formula. If the indices matched exactly, the anchor was counted as correct. The fraction of matched anchors was calculated as:

$$\text{Fraction of Matched Anchors} = \frac{\text{Number of matched anchors}}{\text{Total number of anchors}} \tag{2}$$

### ProteinMPNN Integration

We used ProteinMPNN [11] to sample new peptide binders for a given pMHC backbone structure. In ProteinMPNN, the encoder receives the structure as input rather than the sequence. The decoder can generate sequences either by conditioning on both the structure and existing sequence information (conditional mode) or by using only the encoded backbone (unconditional mode). We applied the conditional mode in this study, as the backbone of MHCs is highly conserved and does not provide sufficient information for peptide sampling across different alleles.

In conditional mode, we distinguish two types of amino acid positions in the structure:

- **Unfixed (designable or variable)**: Amino acid positions whose identities are not provided to the model; the decoder samples new residues for these sites.
- **Fixed (non-designable or conserved)**: Amino acid positions whose identities are provided to the model as conditioning information (along with the backbone) and are therefore not altered during sequence generation.

In PMGen, the MHC sequence is always fixed while the peptide sequence is sampled. By default, only the peptide's anchor positions are fixed. The fraction of randomly fixed non-anchor peptide positions can also be adjusted by the user to control the level of design flexibility.

### NetMHCpan Integration

In the current pipeline, we used NetMHCpan 4.1 [6] for MHC-I and NetMHCIIpan 4.3 [12] for MHC-II. NetMHCpan is applied twice during the PMGen workflow: (i) for anchor prediction prior to PANDORA homology modeling, and (ii) after ProteinMPNN peptide generation to rank candidate peptides and select binders based on their predicted binding affinity and EL percentile rank.

Since NetMHCpan requires an HLA allele and peptide sequence as input, PMGen first searches for compatible alleles using a sequence alignment against known HLA alleles. Once a matching allele is identified, PMGen checks whether it is supported by NetMHCpan. If not, the most similar NetMHCpan-accepted allele is selected based on sequence similarity.

The PMGen pipeline supports two modes of anchor prediction:

1. **Multiple-anchor prediction**: NetMHCpan is run across varying peptide lengths (if the peptide is longer than 8 amino acids). Predicted anchor positions are ranked by EL percentile rank and, secondarily, by predicted binding affinity. The top $k$ anchors (where $k$ is user-defined) are then selected. For each selected anchor, PMGen predicts a distinct peptide–MHC structure. The best AlphaFold-predicted structure is chosen based on its average peptide-pLDDT score.
2. **Single most-reliable anchor**: Only the highest-ranked anchor predicted by NetMHCpan is used, and a single structure is predicted for the peptide–MHC pair.

In the benchmarking experiments reported in this paper, the second mode was used for PMGen initial guess and PMGen+TE, while for PMGen+pLDDT we tested all possible anchor combinations and selected the best based on pLDDT.

### Peptide Generation Analysis

The PDB structures predicted by PMGen were categorized into two groups based on their structural accuracy: high-quality predictions ($n = 55$, pRMSD $< 0.6$) and low-quality predictions ($n = 51$, pRMSD $> 1.0$). Three mutation screens were designed, in which one (screen 1), two (screen 2), or three (screen 3) amino acid positions were treated as variable, while the remaining residues were fixed. The variable residues were defined as contiguous linear segments within the peptide sequence.

For each structure and each screen, ten peptide variants were sampled using ProteinMPNN. Their eluted ligand percentile ranks were predicted using NetMHCpan, and the top three peptides with the lowest ranks were selected. As a control, all possible $20^k \times \mathrm{L}_{\mathrm{peptide}}$ random mutations for screen $k$ were generated. Since the number of random mutations was prohibitively large, we randomly selected 1% of peptides. We ensured that all mutation windows and all screens across all structures and PDB ids have relatively equal fraction of samples. In total, we analyzed 29,802 low pRMSD (good predictions) and 27,161 high pRMSD (bad predictions) cases along with ProteinMPNN-sampled peptides. We then calculated their pRMSD versus the original peptide's predicted structure.

Finally, for each mutation window, enrichment analyses were performed comparing pRMSD distributions of sampled peptides against those of random peptides. The area under the curve (AUC) was computed for each mutation window and subsequently averaged across all windows corresponding to each pMHC structure (Figure S5).

### Fine-tuning ProteinMPNN

**Data preparation** ProteinMPNN is the most widely used model for protein design tasks. However, similar to AlphaFold, it lacks sufficient observations of stable peptide–MHC structures in its training data. In the current study, we used the PMGen pipeline to generate high-quality pMHC structures to fine-tune ProteinMPNN. For this purpose, we first downloaded eluted ligand and binding affinity data from IEDB. To reduce training noise arising from anchor complexity, we used only MHC-I data for training; however, the same methodology is also applicable to MHC-II. We removed all pMHC pairs labeled as non-binders ($\mathrm{EL}_{label} = 0$ or BA $> 500\,\mathrm{nM}$).

To account for MHC diversity, we counted the number of times each MHC allele was observed in combination with different peptides. We then calculated the median of these counts across all sampled MHC alleles and randomly sampled that number of pMHC pairs for each allele, iteratively. All MHC alleles with counts below the median were fully sampled and removed from the next sampling iteration. We continued this procedure with a sampling cap of 1000 samples per allele, meaning that once an allele reached 1000 samples, it was automatically excluded from subsequent iterations. This method ensured that MHC alleles with fewer samples were fully represented, while highly abundant alleles were capped, allowing us to construct a well-distributed MHC dataset. In the next step, we aimed to increase peptide sequence diversity while reducing the dataset size by removing redundant peptides. For this purpose, we assumed anchor positions to be the most conserved residues in a peptide and, to avoid biasing the training toward learning the same anchors for a given MHC, we prioritized anchor diversity. Accordingly, across the sampled dataset, we approximately defined the first and second anchor positions for each peptide-MHC pair. Then, we calculated the frequency of each amino acid at the first and second anchor positions, independently. We applied the same iterative median sampling strategy with an amino acid cap of 1000 for each of anchor residues.

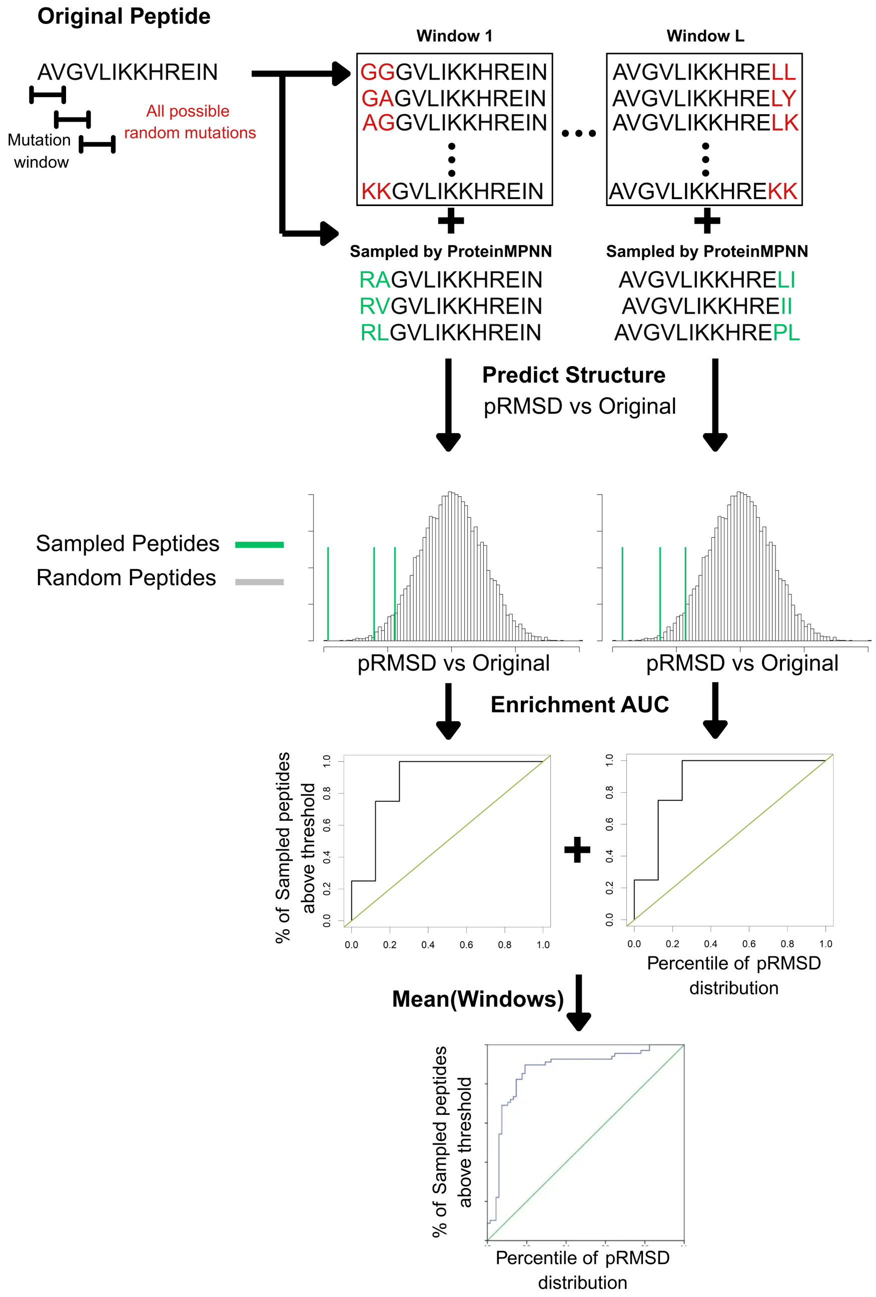


Figure S5: **Mutation screen analysis pipeline.**

We then predicted the structures of the resulting samples using the PMGen initial guess and excluded samples with peptide pLDDT below 80 or peptide PAE above 6.0. In the end, a dataset of 10,216 samples remained and was randomly split into $80\%$ training and $10\%$ validation and test sets each (Figure S6A-B).

**Training** We did not modify the architecture of the ProteinMPNN model by adding or removing any layers. Fine-tuning was performed using the same objective function as in the original method, namely the autoregressive prediction of masked amino acids with a random decoding order. During training, all MHC residues were fixed, while the peptide residues were masked. Consequently, the model was required to predict the correct peptide tokens in a random decoding order while conditioning on the full structure. The fixed MHC positions were not predicted but remained part of the conditioning context. To minimize deviation from, or unlearning of, the original ProteinMPNN model, we employed two adaptation strategies: elastic weight consolidation (EWC) and low-rank adaptation (LoRA). We additionally explored several hyperparameters, including the learning rate and the number of frozen layers. Training was carried out using a masked negative log-likelihood (categorical cross-entropy) loss. Peptide sequence recovery was used as the primary evaluation metric (Figure S6C). Furthermore, we evaluated the model's ability to recover the full pMHC sequence after fine-tuning and compared it to the original ProteinMPNN model to assess potential unlearning effects. Our results indicated that LoRA, without freezing any layers and using a learning rate of $1e-9$, yielded the optimal configuration. The final model exhibited minor forgetting in full pMHC sequence reconstruction (Figure S6D).

### Main Changes in PANDORA Pipeline

To enable PANDORA to operate robustly in the best-anchor-predicted mode, we modified its built-in NetMHCpan prediction logic. In the original implementation, only the best-aligned allele was used for anchor prediction. In our modified version, the top 20 aligned alleles are selected and ranked by alignment score. NetMHCpan is then executed iteratively, starting from the highest-ranked MHC sequence. If the NetMHCpan prediction is successful (i.e., the allele is present in the NetMHCpan's `allelelist.txt` file), the loop terminates and that prediction is used to determine the anchor positions.

This modification improves anchor assignment for MHC queries with low sequence similarity to known alleles, as it allows the algorithm to fall back to progressively less similar alleles until a valid prediction is found.

Additionally, we made minor changes to the `Modeling_Functions.py`, `PMHC.py`, and main `PANDORA.py` scripts to:

- fix bugs affecting NetMHCpan execution,
- modify output parsing for the PMGen pipeline, and
- save predicted anchors and alignment results as `.json` files.

Further details on these modifications are available on our GitHub repository:
`AFfine_PANDORA_modifications.txt`.

### Details on Modeling Software and Prediction Time

PMGen integrates multiple external tools into a single automated pipeline, including NetMHCpan, PANDORA, AlphaFold, and ProteinMPNN. NetMHCpan can be omitted if anchor positions are provided by the user. The core PMGen pipeline is implemented primarily in Python 3.9 and Bash.

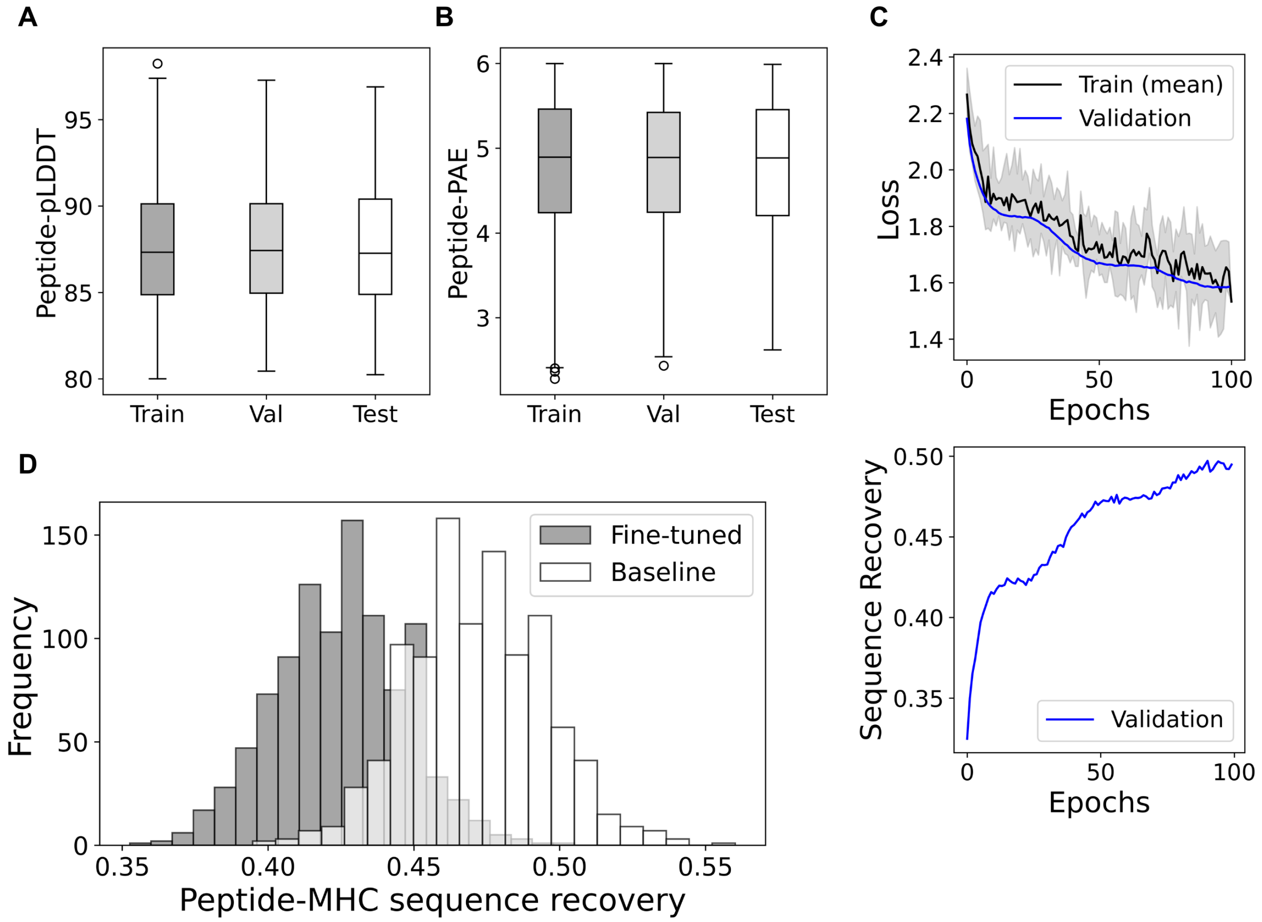


Figure S6: **ProteinMPNN fine-tuning with PMGen structures.** (A-B) Distribution of peptide-pLDDT and PAE for train, validation and test datasets. (C) Training and validation loss (left) and validation sequence recovery (right) monitored over 100 epochs. (D) Sequence recovery of full peptide-MHC complex before and after fine tuning. (E) Peptide sequence recovery before and after fine-tuning.

PMGen does not use multiple sequence alignments (MSAs) for AlphaFold predictions, as omitting MSAs substantially reduces inference prediction time. Furthermore, modifications to the AlphaFold2 pipeline [2] reduce PMGen's software requirements compared to the original AlphaFold2 implementation. PMGen can be executed on both CPU and GPU hardware. Within the pipeline, PANDORA and the Template Engineering step run exclusively on CPU, whereas AlphaFold and ProteinMPNN can run on either CPU or GPU.

**Speed** Using default settings, generation of four engineered templates via Template Engineering takes approximately 20 seconds on a single CPU core. A single AlphaFold prediction in PMGen for an average-sized pMHC complex takes approximately 9 seconds on an NVIDIA A100 GPU and 180 seconds on CPU. For the Initial Guess mode, no template engineering step is performed, and therefore its runtime is not included in this estimate. ProteinMPNN sequence generation for 10 peptides on the same GPU requires roughly 3 seconds.

The reported runtimes do not include the model compilation times for AlphaFold and ProteinMPNN, which are typically around 30 seconds.
